# The effect of heavy metals on thiocyanate biodegradation by an autotrophic microbial consortium enriched from mine tailings

**DOI:** 10.1101/2020.06.13.149401

**Authors:** Farhad Shafiei, Mathew P. Watts, Lukas Pajank, John W. Moreau

## Abstract

Bioremediation systems represent an environmentally sustainable approach to degrading industrially-generated thiocyanate (SCN^-^), with low energy demand and operational costs, and high efficiency and substrate specificity. However, heavy metals present in mine tailings effluent may hamper process efficiency by poisoning thiocyanate-degrading microbial consortia. Here we experimentally tested the tolerance of an autotrophic SCN^-^-degrading bacterial consortium enriched from gold mine tailings for Zn, Cu, Ni, Cr, and As. All of the selected metals inhibited SCN^-^ biodegradation to different extents, depending on concentration. At pH of 7.8 and 30°C, complete inhibition of SCN^-^ biodegradation by Zn, Cu, Ni, and Cr occurred at 20, 5, 10, and 6 mg L^-1^, respectively. Lower concentrations of these metals decreased the rate of SCN^-^ biodegradation, with relatively long lag times. Interestingly, the microbial consortium tolerated As even at 500 mg L^-1^, although both the rate and extent of SCN^-^ biodegradation were affected. This study highlights the importance of considering metal co-contamination in bioreactor design and operation for SCN^-^ bioremediation at mine sites.

**Key points:**

- Both the efficiency and rate of SCN^-^ biodegradation were inhibited by heavy metals, to different degrees depending on type and concentration of metal
- The autotrophic microbial consortium was capable of tolerating high levels of As

## Introduction

For higher organisms, thiocyanate (SCN^-^) is a known goitrogen, i.e. a chemical with deleterious anti-thyroid effects with prolonged exposure (Gaitan 1989); acute SCN^-^ poisoning can also occur (Gould et al. 2012). The toxicity of this compound occurs at blood serum levels greater than 1 mg per 100 mL (Lage et al. 1994). Both chronic and acute toxicity of SCN^-^ to aquatic organisms, including *Daphnia magna* (Parkhurst et al. 1979) and various fish species (e.g., (Bhunia et al. 2000; Kevan and Dixon 1991; Lanno and Dixon 1996)), has also been demonstrated.

Gold processing commonly involves mixing finely-ground ores with the lixiviant sodium cyanide (Woffenden et al. 2008). Gold-bearing ores also naturally contain sulfide minerals that release reduced sulfur species during ore processing, which react with cyanide (CN^-^) to generate SCN^-^ (Mudder et al. 2001). This process can elevate levels of SCN^-^ to higher than 1000 mg L^-1^ in tailings wastewaters (Given and Meyer 1998). Finely ground and chemically processed ore materials and associated wastewaters are typically stored in tailings storage facilities (TSFs) intended to limit the environmental impact and facilitate water-reuse by mining companies (ERR 2007). However, tailings water seepage from TSFs can potentially contaminate the underlying groundwater with SCN^-^ (Kossoff et al. 2014), and sometimes results in elevated levels of this contaminant in groundwater near mine sites (Bakatula et al. 2012).

The conversion of CN^-^ into SCN^-^ is generally promoted in gold ore processing effluent because of the lower toxicity of the latter compound (Bhunia et al. 2000; Ingles and Scott 1987). However, although SCN^-^ is almost seven times less toxic than CN^-^ (ICMI 2012; Woffenden et al. 2008), the greater chemical stability of SCN^-^ compared to its parent compound (Akcil 2003) leads to its accumulation in mining waste streams (Woffenden et al. 2008), as well as its environmental persistence (Mediavilla et al. 2019). Therefore, although not explicitly addressed in regulatory guidelines for discharge of CN^-^-bearing mine effluents, SCN^-^ is still considered by regulatory agencies to be a threat to aquatic wildlife (Bhunia et al. 2000; Gould et al. 2012).

Comprehensive reviews have summarized current chemical and biological treatment methods for either CN^-^ degradation (Gould et al. 2012) or concomitant degradation of CN^-^ and SCN^-^ (Akcil 2003; Botz et al. 2016; Mudder et al. 2001). Compared to physical or chemical approaches, bioremediation systems are considered to be more environmental friendly, efficient (Akcil 2003), cost-effective (Akcil and Mudder 2003; Nelson et al. 1998), and substrate specific (Das and Dash 2014). Accordingly, they constitute a preferred treatment approach in the mining industry, especially when cleaner effluents are targeted (Akcil 2003).

In bioremediation systems for SCN^-^, growth of specific microorganisms is promoted (Watts and Moreau 2018), particularly bacteria that can utilize SCN^-^ as a source of electrons. Some of these microbes also rely only on nitrogen and/or sulfur obtained from SCN^-^ as growth nutrients (Gould et al. 2012). However, tailings effluents typically contain a range of contaminants, including cyanide, cyanate (CNO^-^), ammonia, nitrate, sulfate, chloride, and heavy metals (Given and Meyer 1998; Woffenden et al. 2008) that may inhibit the efficiency of SCN^-^ biodegradation (Gould et al. 2012).

Like most other organisms, microorganisms are susceptible to high concentrations of heavy metals (Giller et al. 1998; Mattila et al. 2007). The effects of heavy metals on biodegradation of environmental pollutants has been studied, mostly for separate hydrocarbons (e.g., (Ma et al. 2018)), mixtures of hydrocarbons (Amatya et al. 2006), diesel oil (Sprocati et al. 2012), polychlorinated biphenyls (PCBs), perchloroethene (PCE) (Lu et al. 2020), and phenanthrene (Wong et al. 2005). The effects of operational conditions such as pH, temperature, loading, phosphate amendment, and light, on SCN^-^ biodegradation, have also been investigated (Kantor et al. 2017; Kantor et al. 2015; Lay-Son and Drakides 2008; Watts et al. 2017a; Watts et al. 2019). Furthermore, the inhibitory effect of NH_4_^+^, another co-contaminant from gold cyanidation (Given and Meyer 1998), on SCN^-^ biodegradation has been studied for pure cultures and microbial consortia in shake flask experiments and laboratory-scale activated sludge bioreactors (Lay-Son and Drakides 2008). However, a lack of knowledge exists on the effects of toxic heavy metals on microorganisms actively degrading SCN^-^.

The present study aimed to evaluate the heavy metals tolerance of a previously characterized autotrophic SCN^-^-degrading consortium enriched from surface tailings of a gold mine in western Victoria, Australia (Watts et al. 2017b). Here, we hypothesized that heavy metals at varying concentrations, associated with tailings effluent, would differentially inhibit the performance of this SCN^-^ biodegrading consortium in shake flask experiments.

## Materials and methods

### Consortium growth conditions

A previously characterized natural autotrophic SCN^-^-degrading consortium (Watts et al. 2017a; Watts et al. 2017b; Watts et al. 2019) was used for this study. Prior to the start of experiments, at least five transfers were performed aseptically with 10% (v/v) of early-stationary phase culture to a 500 mL Erlenmeyer flask containing 180 mL of fresh medium. All cultures were incubated at 30°C and 120 rpm continuous rotation. The medium (1 L) was comprised of Na_2_SO_4_ (2.25 g), NaHCO_3_ (0.25 g), MgSO_4_ (0.51 g), CaCl_2_.2H_2_O (1.25 g), KCl (0.1 g), NaCl (1.5 g), Na_2_HPO_4_.7H_2_O (0.05 g), KSCN (1 g), and 4-(2-hydroxyethyl)-1-piperazineethanesulfonic acid (HEPES) (3.5 g) in ultrapure water (Milli-Q^®^). Micronutrients were supplied through addition of 0.05% (v/v) of Mineral Elixir solution to the medium. The composition of this solution per liter was nitrilotriacetic acid (NTA; free acid non-trisodium salt, 2.14 g), MnCl_2_.4H_2_O (0.1 g), FeSO_4_.7H_2_O (0.3 g), CoCl_2_.6H_2_O (0.17 g), ZnSO_4_.7H_2_O (0.2 g), CuCl_2_.2H_2_O (0.03 g), AlK(SO_4_)_2_.12H_2_O (0.005 g), H_3_BO_3_ (0.005 g), Na_2_MoO_4_.2H_2_O (0.09 g), NiSO_4_.6H_2_O (0.11 g), and Na_2_WO_4_.2H_2_O (0.02 g). After pH adjustment at 7.5±0.02 using 1 N NaOH and 1 N HCl, the medium was autoclaved at 121°C for 15 min.

### Inoculum preparation

For all experiments, a 36-h culture was centrifuged (Eppendorf Centrifuge 5810) at 5000 rcf for 10 min at room temperature. Cells were washed twice with a half-volume of sterile saline solution (pH 7.0±0.6), and pellets were resuspended in fresh medium before inoculation. Optical density at 600 nm wavelength (OD_600_) was measured to monitor cell growth.

### Metal solutions

Stock solutions of zinc (as Zn^II^), copper (as Cu^II^), chromium (as Cr^VI^), nickel (as Ni^II^), and arsenic (as As^V^) were prepared immediately before use by dissolving ZnCl_2_, CuCl_2_.2H_2_O, K_2_CrO_4_, NiCl_2_.6H_2_O, and Na_2_HAsO_4_.7H_2_O in ultrapure water (Milli-Q^®^), respectively. Stock solutions were further diluted and filtered via 0.22 μm syringe filters (Millex^®^ Express PES Membrane). These metals were selected based on peer-reviewed literature (Ebbs et al. 2010; Fashola et al. 2016; Noble et al. 2010) and proprietary data on metals concentrations in tailings effluent from an operating Victorian gold mine. Background Ni, Zn and Cu concentrations derived from medium reagents were all <1 mg L^-1^ (data not shown).

### Metal tolerance experiment

Prior to each experiment, all glassware was soaked in nitric acid (2.5% for 24 hours) and rinsed with ultrapure water to remove any trace metals. Four concentrations were used for each heavy metal: 10, 20, 40, and 60 mg L^-1^ for Zn^2+^; 0.5, 1.5, 2.5, and 5 mg L^-1^ for Cu^2+^; 5, 7.5, 10, and 15 mg L^-1^ for Ni^2+^; 1.5, 3, 6, and 30 mg L^-1^ for Cr^6+^; and 10, 30, 300, and 500 mg L^-1^ for As^5+^. Previous toxicity studies of these metals with different groups of microorganisms were consulted to determine experimental concentrations (Alexandrino et al. 2011; Chiboub et al. 2016; Ma et al. 2019; Takeuchi et al. 2007). For each metal, 5% (v/v) pre-prepared metal solution was added to 250 mL of sterile culture medium. Filtered ultrapure water (no metals) was used for metal-free (i.e., positive) controls. All flasks were inoculated with 10% (v/v) of cell suspension. For negative controls, the inoculum was replaced with the same volume of cell-free culture medium. Furthermore, killed-cell controls were used for metal-free trials to ensure that the observed SCN^-^ degradation is a result of cell metabolism. Cu experiments, however, included killed-cell controls for all metal amendment levels as well. These extra controls were used to investigate whether the drop in dissolved Cu throughout the experiments was due to cell adsorption or metabolic activity. To prepare killed-cell controls, washed cells were resuspended in fresh culture medium followed by autoclaving. All cultures were incubated in triplicate simultaneously for one type of metal at a time. Statistical analyses were performed using Minitab^®^ 18.1 (Minitab 18 Statistical Software 2017).

Initial and final experimental SCN^-^ concentrations (mg L^-1^) were used in Equation (1) to calculate SCN^-^ biodegradation efficiency (%). Lag time was defined as hours with insignificant change to SCN^-^ level, as determined by dependent t-test (P ≤ 0.05). Equation (2) was used to calculate SCN^-^ biodegradation rate. The duration of biodegradation was defined as either 5 days (i.e., time from beginning to end of the experiment), or the time until SCN^-^ levels remained constant.

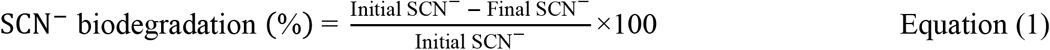

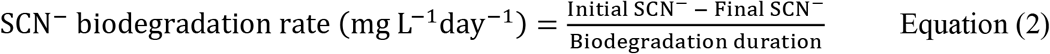

### Thiocyanate measurement

Shake flasks were sampled at regular intervals over the five days of incubation. For SCN^-^ measurements, less than 0.8 mL of medium was centrifuged at 16,000 rcf for 10 min at room temperature to pellet cells. The supernatant was removed and stored at −20°C until quantification. A colorimetric method was adapted to determine SCN^-^ concentration (Baird et al. 2017). For this analysis, each sample was mixed with ferric nitrate reagent, and absorbance was measured at 460 nm wavelength.

### Metal analyses

Sampling for dissolved metals was performed in the same way as SCN^-^ analyses, except that the supernatant was stored at 4°C prior to analysis. Furthermore, at sampling intervals, less than 2.5 mL of the medium was centrifuged at 5,000 rcf for 10 min, and the pH of the supernatant was measured.

Sample aliquots of between 10 and 60 microliters were diluted with 0.4 N double-distilled nitric acid solution containing 10 ppb Rh as an internal standard. Single element standards spiked with 10 ppb Rh were used for calibration. Sample dilution factors varied from 50 to 300, depending on the original analyte concentration. Sample solutions were then vortexed briefly, prior to analysis on an Agilent 7700x ICP-MS in collision cell mode using 3 ml per minute He as the carrier gas.

### Geochemical modeling

Metal speciation under experimental conditions was calculated using PHREEQC (Parkhurst and Appelo 2013). A modified Minteq (version 4) database (U.S. Environmental Protection Agency 1998) was used for all simulations, with pH 7.7 and a temperature of 30°C as input parameters.

## Results

### Measured dissolved metals concentrations by ICP-MS

Dissolved Zn and Cu concentrations over time for Zn- and Cu-amended experiments are shown in Supplementary Figure 1. These metals were the only ones observed to change in concentration over time, e.g., dissolved Zn concentrations decreased significantly over the first 40 hours of incubation. In Cu-amended experiments, aqueous Cu concentrations generally decreased in inoculated cultures over the 120-hour incubation period, with the exception of an apparent increase from an initially lower-than-added value (by nearly half) for the highest Cu level (added as 5 mg L^-1^). All other metal amendments showed no significant changes over the incubation period for any initial concentration (Supplementary Figure 2).

### pH monitoring

Previous studies have reported the effect of pH on metal toxicity to microorganisms, e.g., (Moberly et al. 2010; Van Nostrand et al. 2005). Therefore, the pH of cultures was monitored across sampling timepoints. The initial addition of filtered heavy metal solutions to the culture medium did not change pH significantly; neither was any difference observed between initial pH values for any experimental conditions. Average pH values were 7.75 ± 0.03 and 7.73 ± 0.06 for the start and end, respectively, of all experiments for all trials and controls.

### Metal speciation calculations

Generally, increasing levels of metals in culture amendments produced increasing dissolved metals concentrations, as well as higher saturation indices (SI) with respect to respective mineral phases. PHREEQC modeling predicted that almost all Zn in the culture medium was present as dissolved sulfide species (Table S1), with positive or close to zero SI values with respect to sphalerite, wurtzite, and amorphous ZnS (Table S2). Likewise, with aqueous Cu(HS)_3_^-^ as the predominant dissolved Cu species (Table S3), Cu was predicted to be near or at saturation with respect to copper sulfides (Table S4).

In metal-free experiments (i.e., positive controls), approximately 99.6% of Ni (from medium ingredients) was present as Ni(NTA)^-^ (Table S5). The next most prevalent Ni species in these experiments were Ni(NTA)_2_^4-^, Ni^2+^, and NiOH(NTA)^2-^, each accounting for only about 0.1 percent of total Ni. Total Ni at amendment levels of 5, 7.5, 10, and 15 mg L^-1^ was comprised of about 64%, 43%, 32%, and 22% Ni(NTA)^-^ and about 24%, 38%, 45%, and 53% Ni^2+^, respectively. Relatively large positive SI values were calculated for nickel sulfide phases (Table S6), as well as for Ni-molybdate (at 7.5, 10, and 15 mg L^-1^ Ni) and Ni-phosphate (at 15 mg L^-1^ Ni).

Modeling of Cr-amended experiments predicted that most added Cr was present as chromate (Table S7). None of the predicted Cr phases showed negative or close to zero SI values (Table S8). Four arsenate species were predicted in As-amended experiments (Table S9), including HAsO_4_^2-^ as 94-95% of total As. Like Cr, As was largely undersaturated with respect to any phase (Table S10).

### Heavy metals influences on SCN^-^ biodegradation

The influence of heavy metals on SCN^-^ biodegradation is summarized in Table 1 in terms of SCN^-^ biodegradation efficiency (%), rate, and lag time for all inoculated experiments. None of the negative and killed-cell controls showed SCN^-^ biodegradation.

**Table 1.** The influence of heavy metals amendment on SCN^-^ biodegradation

| Heavy metal | Amendment level<br>(mg L<sup>-1</sup>) | SCN<sup>-</sup> biodegradation |  |  |
| --- | --- | --- | --- | --- |
|  |  | Extent (%) | Rate (mg L<sup>-1</sup> day<sup>-1</sup>) | Lag time (h) |
| Zn | 0 | 100 | 377.1 | 6 |
|  | 10 | 100 | 143.0 | 48 |
|  | 20 | 0 | 0 | NA |
|  | 40 | 0 | 0 | NA |
|  | 60 | 0 | 0 | NA |
| Cu | 0 | 100 | 384.2 | < 6 |
|  | 0.5 | 100 | 183.4 | 32 |
|  | 1.5 | 100 | 158.8 | 48 |
|  | 2.5 | 30 | 38.9 | 85 |
|  | 5 | 0 | 0 | NA |
| Ni | 0 | 100 | 379.8 | < 6 |
|  | 5 | 23 | 29.5 | 48 |
|  | 7.5 | 6 | 7.6 | 56 |
|  | 10 | 0 | 0 | NA |
|  | 15 | 0 | 0 | NA |
| Cr | 0 | 100 | 319.4 | 6 |
|  | 1.5 | 100 | 160.6 | 6 |
|  | 3 | 16 | 20.8 | 85 |
|  | 6 | 0 | 0 | NA |
|  | 30 | 0 | 0 | NA |
| As | 0 | 100 | 380.4 | 6 |
|  | 10 | 100 | 392.1 | 6 |
|  | 30 | 100 | 206.2 | 6 |
|  | 300 | 34 | 113.1 | < 6 |
|  | 500 | 31 | 101.6 | < 6 |

NA = Not applicable
< 6: Shorter than the length of time before the first sampling point

### Zinc

The effect of Zn additions on SCN^-^ biodegradation is presented in Fig. 1. At Zn concentrations of 20, 40, and 60 g L^-1^, SCN^-^ biodegradation was completely inhibited. At 10 mg L^-1^ Zn, SCN^-^ was completely degraded within 4.5 days, after a lag period of roughly 48 hours. Furthermore, a slower biodegradation rate (143.0 mg L^-1^ day^-1^) was observed for 10 mg L^-1^ Zn, compared to Zn-free cultures (377.1 mg L^-1^ day^-1^).

**Fig. 1.**
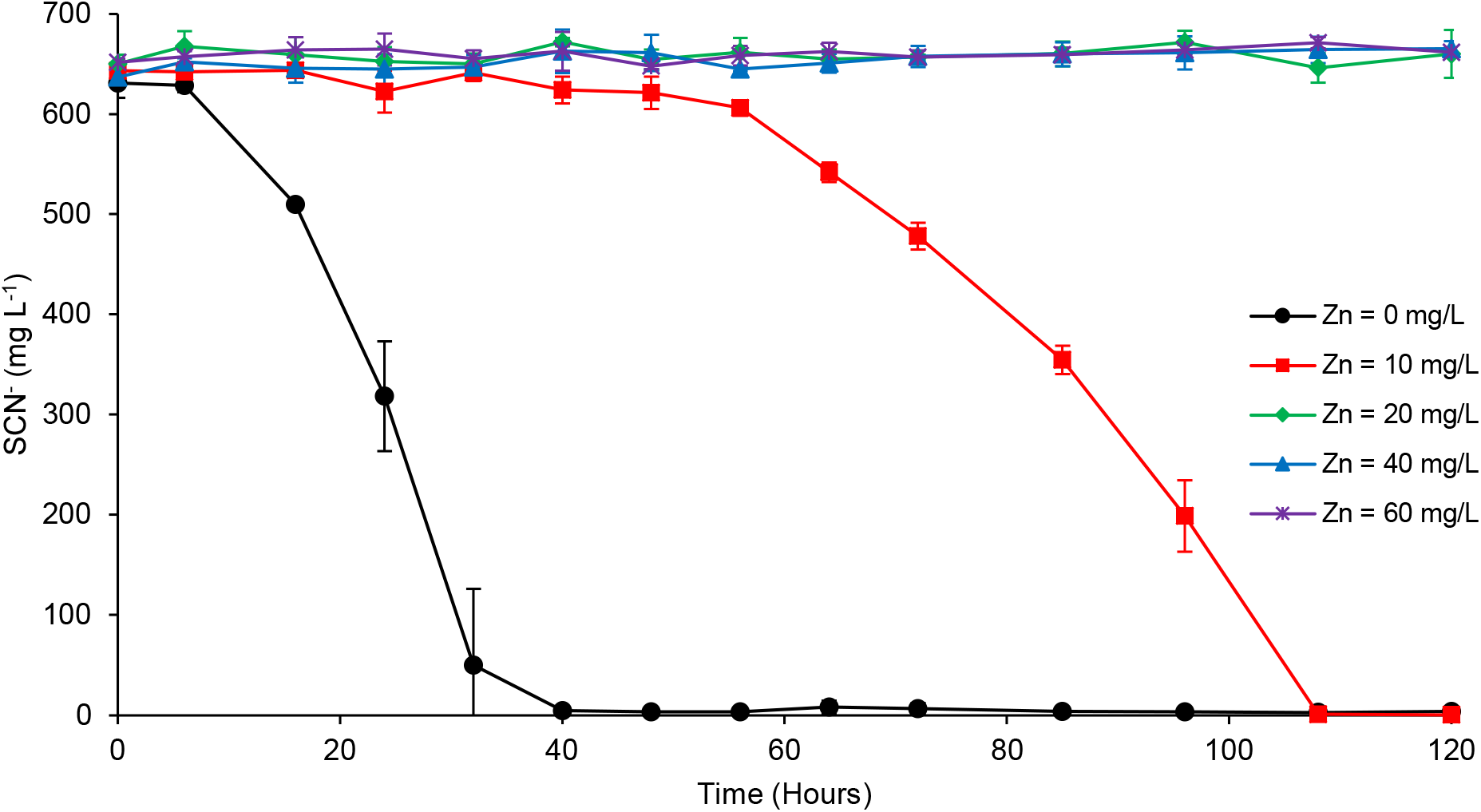
Effect of Zn amendment on microbial SCN^-^ degradation

### Copper

At 5 mg L^-1^ Cu, SCN^-^ biodegradation was completely inhibited (Fig. 2). With 2.5 mg L^-1^ Cu, ~30% SCN^-^ was degraded, with a ~85-hour lag time. Complete SCN^-^ biodegradation was observed at 1.5 and 0.5 mg L^-1^ Cu, with lag times of ~48 and ~32 hours, and biodegradation rates of 158.8 and 183.4 mg L^-1^ day^-1^, respectively.

**Fig. 2.**
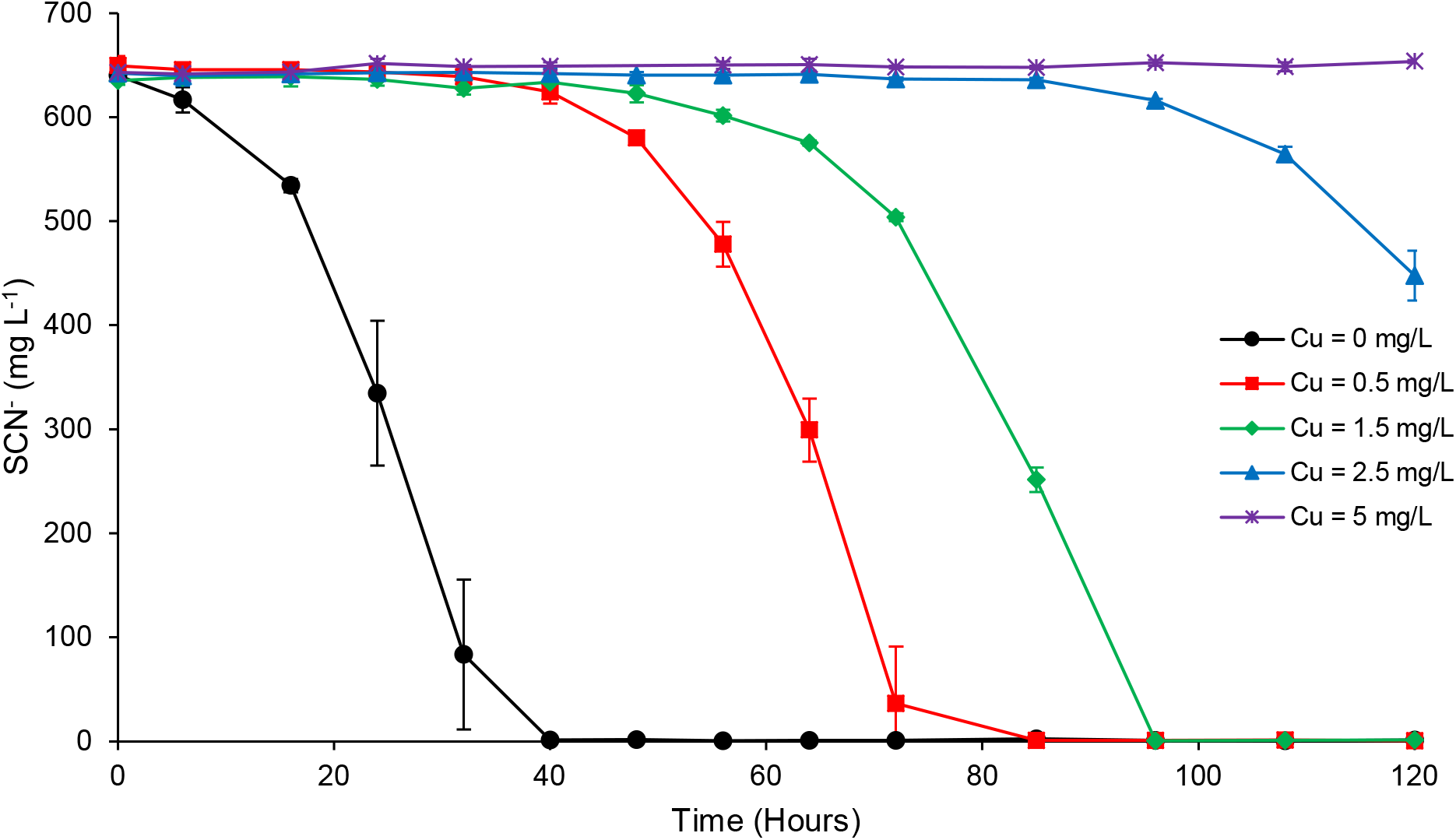
Effect of Cu amendment on microbial SCN^-^ degradation

### Nickel

At 15 and 10 mg L^-1^, nickel completely inhibited SCN^-^ biodegradation (Fig. 3). Only 6% degradation was observed at 7.5 mg L^-1^ Ni, with a ~56-h lag period. At 5 mg L^-1^ Ni, 23% of initial SCN^-^ was degraded after a 48-h lag period.

**Fig. 3.**
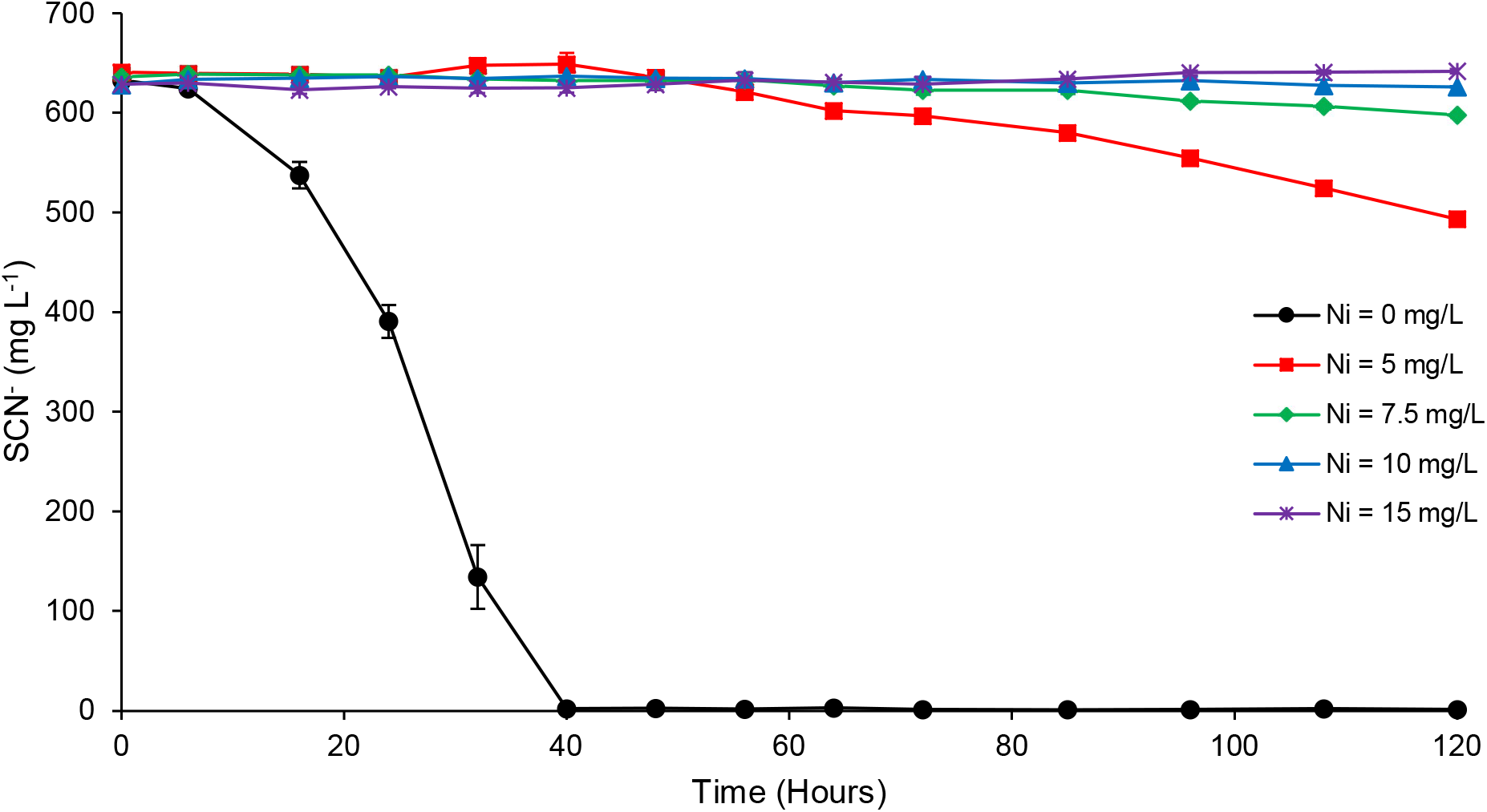
Effect of Ni amendment on microbial SCN^-^ degradation

### Chromium

The effect of Cr amendment on SCN^-^ biodegradation is presented in Fig. 4. At concentrations of 30 and 6 mg L^-1^ Cr, no SCN^-^ was degraded. With 3 mg L^-1^ Cr, only 16% SCN^-^ was degraded over the last 35 hours of the experiment. Amending the culture medium with 1.5 mg L^-1^ Cr did not inhibit complete SCN^-^ biodegradation; however, this level of Cr decreased the degradation rate to 160.6 mg L^-1^ day^-1^, compared to 319.4 mg L^-1^ day^-1^ in Cr-free cultures.

**Fig. 4.**
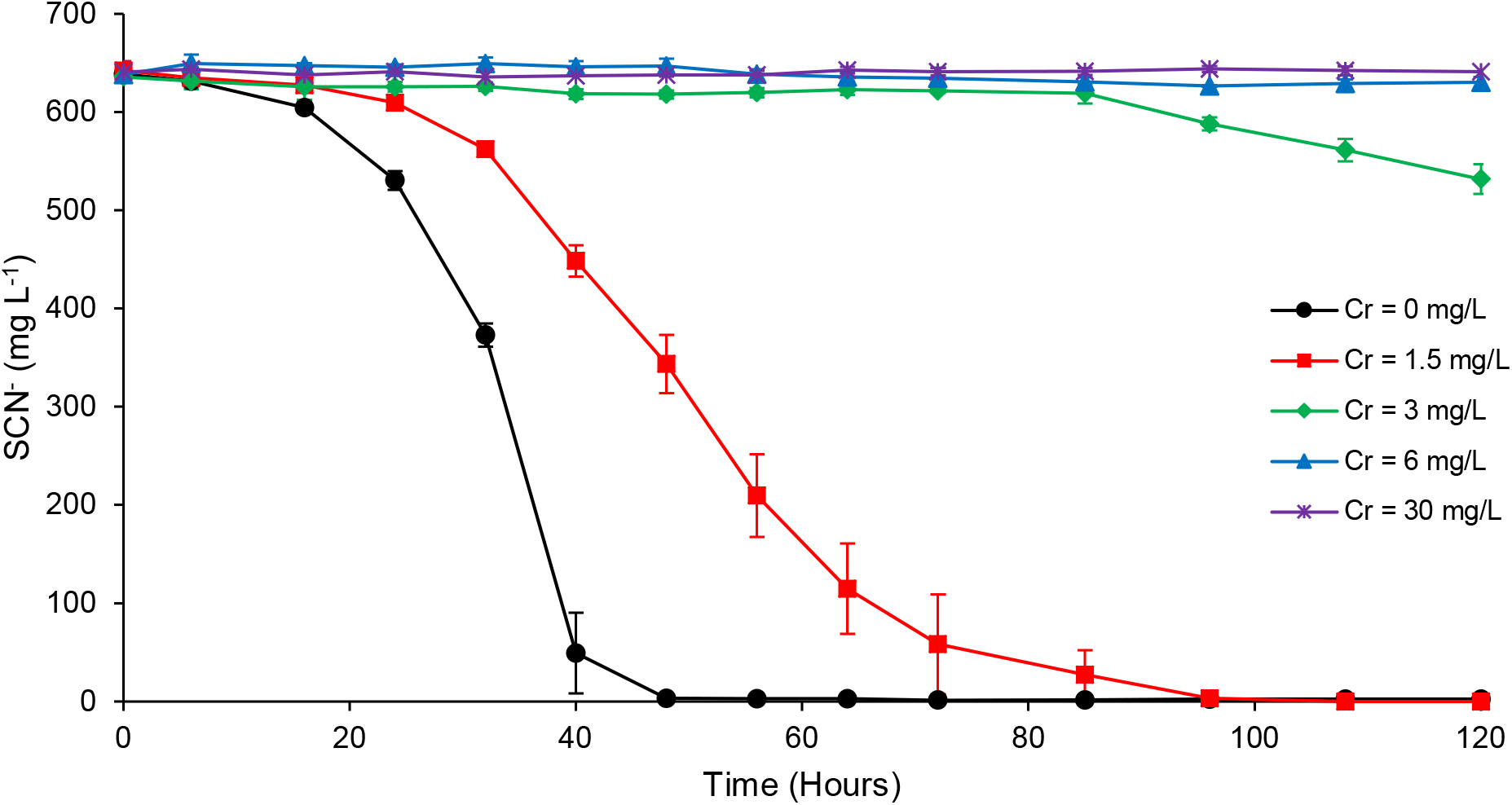
Effect of Cr amendment on microbial SCN^-^ degradation

### Arsenic

The experimental consortium seemed quite tolerant to arsenic even at higher levels of this metalloid. Fig. 5 presents how SCN^-^ biodegradation was affected by varying concentrations of As. Complete SCN^-^ degradation was observed at As concentrations of 10 and 30 mg L^-1^ within 40 and 72 hours of incubation, respectively. These levels of As influenced neither biodegradation extent nor lag time, compared to As-free experiments. A slightly faster biodegradation rate (392.1 mg L^-1^ day^-1^) was calculated for experiments with 10 mg L^-1^ As than for As-free trials (380.4 mg L^-1^ day^-1^), which probably resulted from a slightly higher SCN^-^ concentration in As-amended flasks. Adding 30 mg L^-1^ As to the culture medium led to a biodegradation rate of 206.2 mg L^-1^ day^-1^. The SCN^-^ biodegradation rate in cultures amended with a relatively high concentration of 300 mg L^-1^ As was 113.1 mg L^-1^ day^-1^. At this As level, 34% of SCN^-^ was degraded over the first 48 hours. The same pattern was observed for experiments amended with 500 mg L^-1^ As, with slightly lower biodegradation efficiency (31%) and rate (101.6 mg L^-1^ day^-1^).

**Fig. 5.**
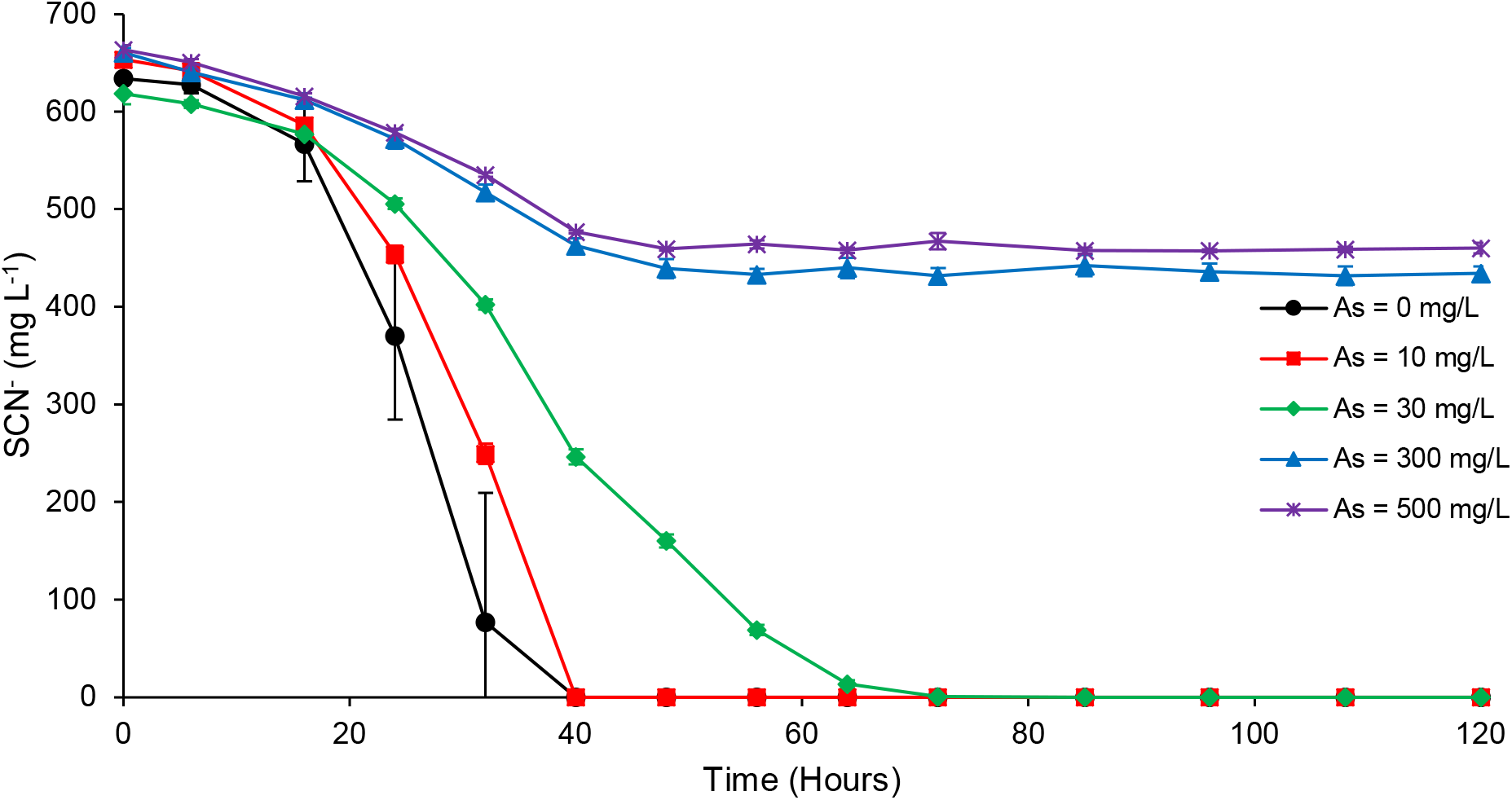
Effect of As amendment on microbial SCN^-^ degradation

## Discussion

Each of the metal(loid)s tested here inhibited SCN^-^ biodegradation to an extent and duration that depended on both type and concentration. Biodegradation of SCN^-^ was fully inhibited at concentrations ≥ 20, 5, 10, and 6 mg L^-1^ for Zn, Cu, Ni, and Cr, respectively, consistent with previous reports for metal toxicity to non-thiocyanate degrading microorganisms (Alexandrino et al. 2011). However, these values represent controlled laboratory experiments, and other experimental or environmental metal toxicity thresholds may differ, depending on geochemical conditions (e.g., (Alexandrino et al. 2011; Cabrera et al. 2006; Sani et al. 2001; Utgikar et al. 2003)).

Lag time for the onset of SCN^-^ biodegradation was a useful measure for comparing metal toxicity effects. For example, while similar biodegradation rates were measured for 1.5 mg L^-1^ each of Cu and Cr, an 8-fold longer lag phase was observed for Cr, indicative of greater toxicity to the microbial consortium from hexavalent Cr. In contrast, the microbial consortium seemed quite tolerant to As over the range of 10 to 500 mg L^-1^; none of the As levels tested in this study extended the lag time for SCN^-^ biodegradation, compared to As-free trials.

The ordering of microbial metals tolerance in this study was As>Zn>Ni>Cu>Cr, in terms of the ability of the consortium to degrade SCN^-^ at similar metal concentrations. Overall, higher tolerance to Zn when compared with Cu was observed in studies using sulfur-oxidizing thiobacilli (Huber and Stetter 1990), iron-oxidizing *Thiobacillus ferrooxidans* (Magnin et al. 1998), and sulfate-reducing bacteria for either single strains (Poulson et al. 1997; Sani et al. 2001) or mixed cultures (Hao et al. 1994; Utgikar et al. 2001; Utgikar et al. 2003). Presumably this observed difference in Zn and Cu toxicity reflected to some degree the relative solubilities of Zn- and Cu-sulfides, which our modeling predicted to be a factor controlling the speciation of these metals in our experiment with SCN^-^ as well.

Lower threshold concentrations for toxicity were previously observed for Ni when compared with Zn under the same experimental conditions (Hao et al. 1994; Huber and Stetter 1990; Poulson et al. 1997), again potentially due to the tendency for Zn to form Zn-sulfides over Ni in mixed metal solutions, as predicted by PHREEQC. Divalent Ni competes catabolically and allosterically with Zn and Fe and therefore interferes with metalloenzyme activities, as well as generates reactive oxygen (Macomber and Hausinger 2016). Therefore, the observed inhibition of SCN^-^ biodegradation in Ni-amended experiments may be attributed to Ni^2+^ species that comprised a larger proportion of total Ni at higher amendment levels, as modeled. The consortium that was used here is dominated by thiobacilli (Watts et al. 2017b; Watts et al. 2019).

The higher toxicity of Cr, when compared with other metals from this study, to SCN^-^ biodegrading microorganisms is in accordance with the results of previous studies; even though with much higher metal concentrations compared to our experiments. An 8-fold higher sensitivity of iron-oxidation by *Thiobacillus ferrooxidans* was reported for Cr when compared with Cu (Magnin et al. 1998). Almost all added Cr in our experiments was predicted to be speciated as chromate, with known adverse effects on bacterial cells (Ramírez-Díaz et al. 2008). We can speculate therefore that the relatively low tolerance of this consortium to Cr is due to the lack or inefficiency of mechanisms required for microbial resistance to chromate (e.g., efflux systems and chromate reductases) (Ramírez-Díaz et al. 2008).

Interestingly, both Cu and Zn concentrations decreased to some degree at the start of incubation periods for inoculated cultures, i.e., positive controls (Suppl. Fig. 1). We hypothesize therefore that these metals partially adsorbed to microbial cells (preferentially to Ni, Cr or As) during the course of our experiments, with Zn adsorbing faster, but to a lesser extent. Alternatively, it is possible that Zn and Cu were incorporated into nanophase Zn- and Cu-sulfides, some of which aggregated to be removed by filtration before ICP-MS analysis. Zn levels decreased asymptotically to a limited extent of removal from solution, while Cu displayed a less systematic trend, decreasing faster than Zn to near complete removal from solution from ~40 hours of incubation for all but the highest Cu concentration (5 mg L^-1^). In any case, we acknowledge that such processes as just described may have impacted on SCN^-^ biodegradation efficiency by effectively reducing metal bioavailability and therefore also metal toxicity. Further work beyond the scope and aim of this study is needed to elucidate such levels of detail and test the above hypotheses. Utgikar et al. (2004) described microbial tolerance for Zn and Cu as a function of metal concentration and exposure time, and we add here the possibility of speciation-driven vital effects. The total concentrations of heavy metals were measured throughout the experiments as filtered (i.e., dissolved) metal levels, and for all the experiments, pH remained constant at 7.7-7.8. PHREEQC modeling predicted conditions exceeding or near to saturation with respect to the precipitation of certain Zn and Cu sulfide phases, as opposed to undersaturated conditions for Ni, Cr and As. Thus, the results of our incubation experiments are consistent with our hypotheses, and exemplify to a degree the links between metal speciation and toxicity. Speculatively, it is also possible, where dissolved Cu levels were observed to increase slightly towards the end of our experiments, that release of Cu from SCN^-^-degrading bacterial cells, in stationary and/or death phases, could explain this observation. Previous findings of the incorporation of Cu in one of the main enzymes involved in autotrophic SCN^-^ biodegradation (Tikhonova et al. 2020), along with the inhibitory effect of cell-associated Cu (De Schamphelaere et al. 2005), would support this speculation.

Arsenic tolerance as observed in the present study was higher than that reported for an endophytic *Citrobacter* strain (400 mg L^-1^) (Selvankumar et al. 2017) or for *Bacillus* isolates (225 and 90 mg L^-1^) (Taran et al. 2019), or *Thiobacillus caprinus* isolates (Huber and Stetter 1990) in similar experiments. For the latter species, for example, a much lower degree of tolerance was observed for As when comparing with Zn, Ni, and Cu (Huber and Stetter 1990). Complete inhibition of SCN^-^ biodegradation did not occur even at the highest As level (500 mg L^-1^), at which approximately 33% of initial SCN^-^ was degraded within the first 48 hours of incubation. However, no further SCN^-^ degradation was observed in As-amended trials beyond this timepoint, highlighting both As concentration and exposure time as a possible considerations for evaluating As toxicity. It is noteworthy that in previous studies, growth inhibition was the primary indicator of As toxicity, rather than the effect of As on a specific phenotypic function, e.g., thiocyanate biodegradation.

The observed tolerance of our experimental consortium for As may indicate an *in situ* selective pressure from the source material used to enrich the experimental microbial consortium. This consortium originated from a gold mine (Watts et al. 2017b) where sulfide ores on average contain substantial levels of As (e.g., 225 mg kg^-1^) (Noble et al. 2010). In fact, arsenopyrite is a main source of As in the mine tailings (0.2-0.4%) (King et al. 2008), and can incorporate As up to 10% w/w (Abraitis et al. 2004). Analysis of soil samples near the mine site from which cultures used for this study were enriched revealed 16-946 mg As (background values: 1-16 mg), 18-740 mg Cr (background: 26-143), and 12-430 mg Pb (background: 9-23 mg) per kg soil (Noble et al. 2010). Chemical monitoring of decant water of the mining site where our consortium was enriched from showed concentration (mg L^-1^) ranges of about 0.03-3.33 for Cu, 0.08-0.61 for As, 0.02-0.26 for Ni and 0.01-0.17 for Zn (personal communication). We note that high levels of As tolerance have been reported previously, e.g., isolates of *Bacillus* sp. and *Aneurinibacillus* sp. that were able to grow in over 1 g L^-1^ As. These isolates were similarly cultivated from As-contaminated (ground)water samples (Dey et al. 2016).

Metal toxicity is determined by both biotic effects and abiotic factors, i.e. physicochemical characteristics of the environment (Babich et al. 1980; Gadd and Griffiths 1977). Microbial inhibition by metals varies with respect to type of microorganism and metal (Babich et al. 1980; Sadler and Trudinger 1967), metal speciation and bioavailability (Wang et al. 2007), and presence of other chemicals such as chelating agents (Sadler and Trudinger 1967) and environmental solutes (Babich et al. 1980). Therefore, higher levels of metals do not necessarily imply greater toxicity (Sadler and Trudinger 1967). Also, higher toxicity has been reported in multi-metal experiments than for single metal assays (Li and Ke 2001a; Li and Ke 2001b; Utgikar et al. 2004), and previous studies have demonstrated greater degrees of metal tolerance of microbial consortia and co-cultures, when compared to isolates (Lu et al. 2020; Ma et al. 2018). This study provides a first step in evaluating the sensitivity of microbial consortia in SCN^-^ bioremediation systems towards potential heavy metal co-contaminants in tailings waste streams. As actual influent to SCN^-^ bioremediation systems likely contains multiple metal co-contaminants, synergistic effects of these metals should also be considered in future research.

## Acknowledgements

The authors wish to acknowledge David Coe, Peter Wemyss and Cameron Hope of Stawell Gold Mine for providing access to mine tailings for enrichment culturing and historical geochemical data. Also, we are grateful to Alan Greig for help with ICP-MS analyses, Christopher Aitken for initial observations that helped to guide experiments, and Angus Keillar for assistance with geochemical modeling.

## Declarations

### Ethics approval

This article does not contain any studies with human participants or animals performed by any of the authors.

### Consent to participate

Not applicable

### Consent for publication

The authors approved the manuscript and gave their consent for its submission to Applied Microbiology and Biotechnology.

### Availability of data and material

Not applicable

### Code availability

Not applicable

### Authors’ contributions

FS and JWM devised the research and wrote the manuscript with critical contributions from MPW. FS conducted all experiments and analyses, with assistance for some tests from LP.

## Supplementary Material

**Supplementary Figure 1.**
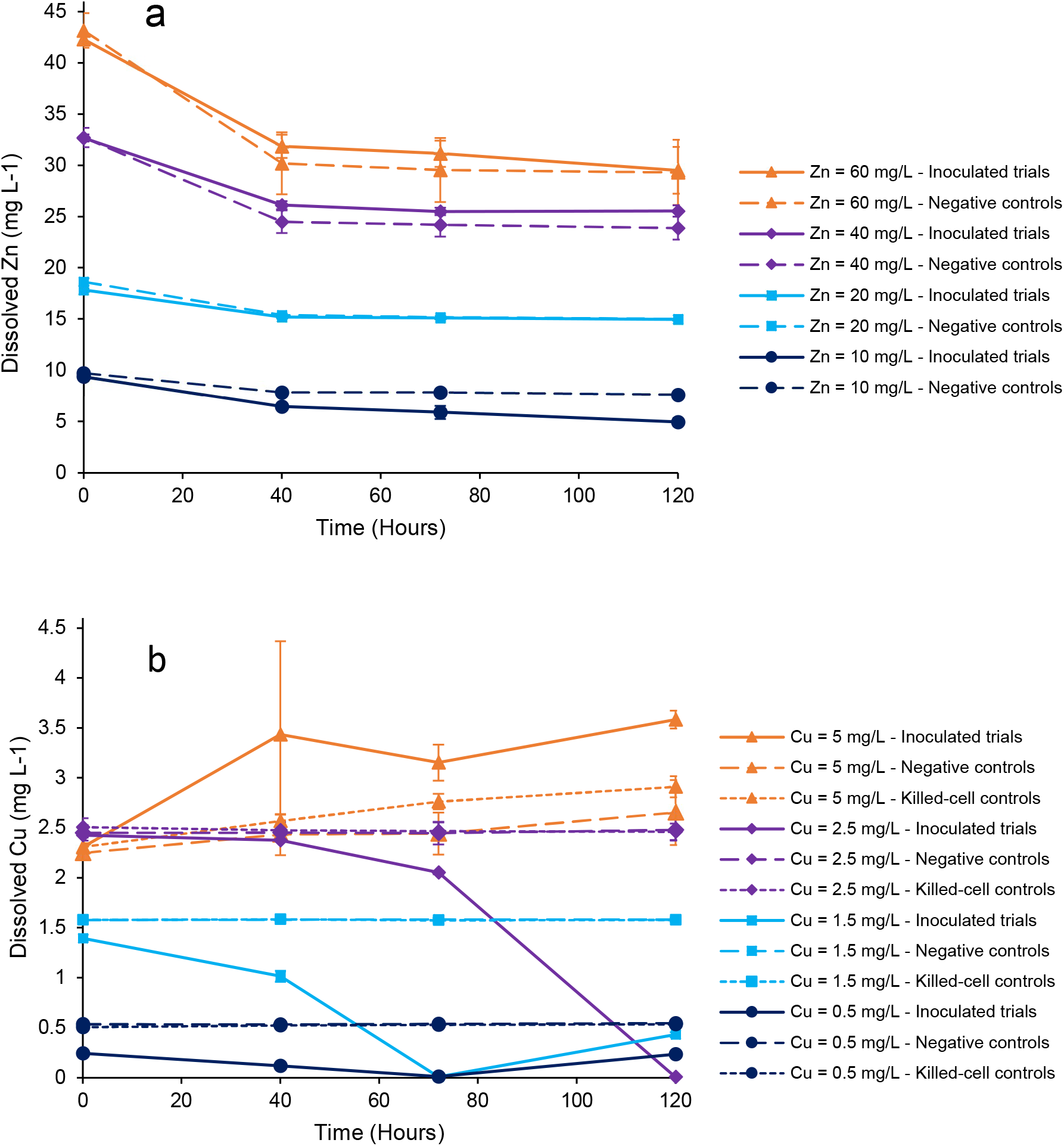
Change in dissolved metals concentrations in metal-amended trials. (**a**) Zn and (**b**) Cu

**Supplementary Figure 2.**
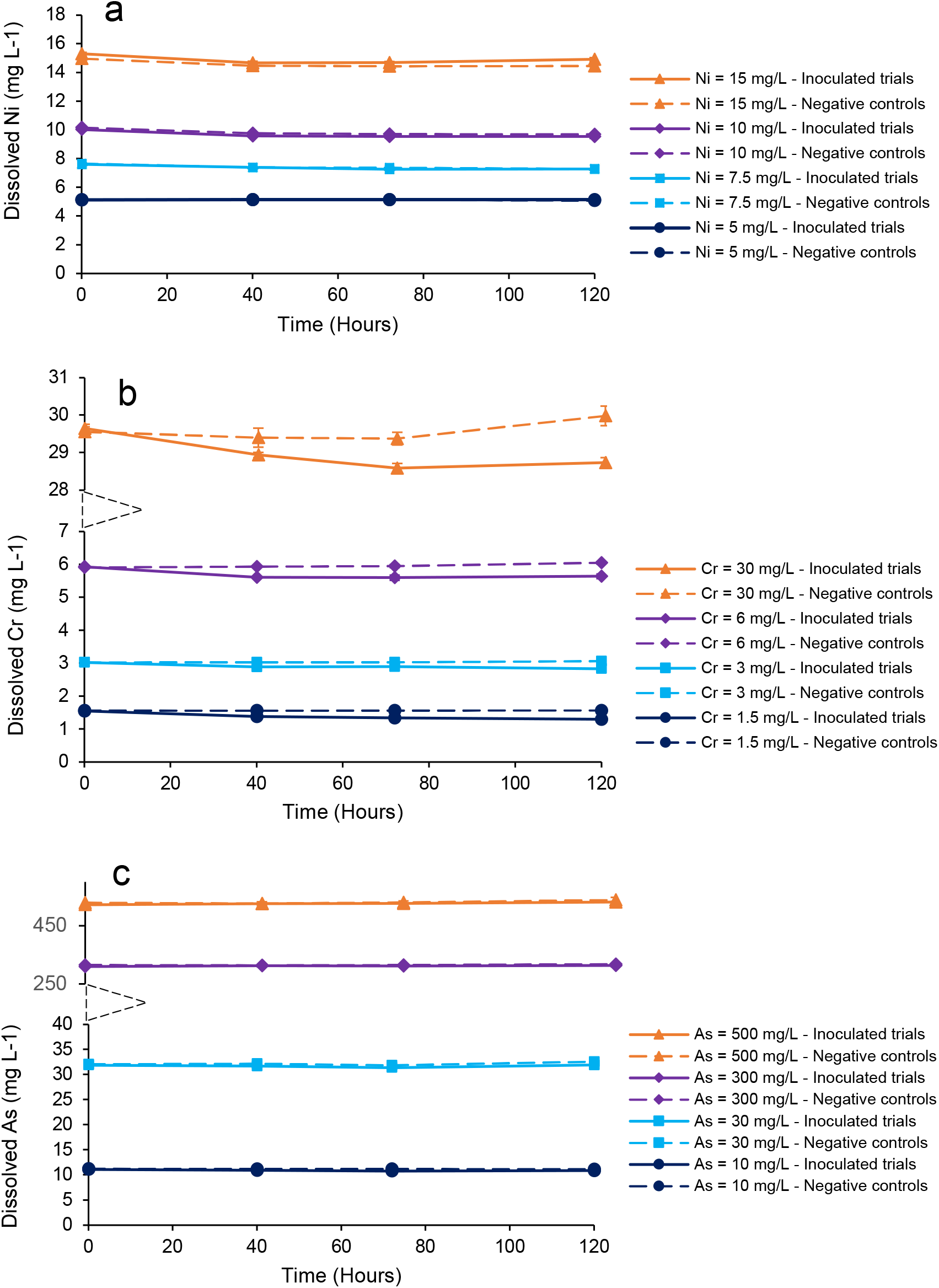
Change in dissolved metals concentrations in metal-amended trials. (**a**) Ni, (**b**) Cr, and (**c**) As

**Table S1.** Distribution (molality) of predicted Zn species at different levels of Zn amendment and non-amended controls

| Zn species | Zn amendment level (mg L<sup>-1</sup>) |  |  |  |  |
| --- | --- | --- | --- | --- | --- |
|  | 0 | 10 | 20 | 40 | 60 |
| ZnS(HS)<sup>-</sup> | $2.83 \times 10^{-6}$ | $1.28 \times 10^{-4}$ | $2.54 \times 10^{-4}$ | $5.12 \times 10^{-4}$ | $7.76 \times 10^{-4}$ |
| Zn(HS)<sub>3</sub><sup>-</sup> | $6.18 \times 10^{-7}$ | $2.71 \times 10^{-5}$ | $5.21 \times 10^{-5}$ | $9.76 \times 10^{-5}$ | $1.37 \times 10^{-4}$ |
| Zn(HS)<sub>2</sub> | $3.95 \times 10^{-8}$ | $1.79 \times 10^{-6}$ | $3.56 \times 10^{-6}$ | $7.16 \times 10^{-6}$ | $1.08 \times 10^{-5}$ |
| ZnS(HS)<sub>2</sub><sup>2-</sup> | $1.01 \times 10^{-8}$ | $4.43 \times 10^{-7}$ | $8.51 \times 10^{-7}$ | $1.60 \times 10^{-6}$ | $2.24 \times 10^{-6}$ |
| Zn(HS)<sub>4</sub><sup>2-</sup> | $3.75 \times 10^{-10}$ | $1.59 \times 10^{-8}$ | $2.96 \times 10^{-8}$ | $5.17 \times 10^{-8}$ | $6.73 \times 10^{-8}$ |
| Zn(NTA)<sup>-</sup> | $7.56 \times 10^{-14}$ | $3.54 \times 10^{-12}$ | $7.53 \times 10^{-12}$ | $1.75 \times 10^{-11}$ | $3.08 \times 10^{-11}$ |
| ZnOH(NTA)<sup>2-</sup> | $5.32 \times 10^{-16}$ | $2.49 \times 10^{-14}$ | $5.30 \times 10^{-14}$ | $1.23 \times 10^{-13}$ | $2.17 \times 10^{-13}$ |
| Zn<sup>2+</sup> | $5.15 \times 10^{-16}$ | $2.49 \times 10^{-14}$ | $5.31 \times 10^{-14}$ | $1.23 \times 10^{-13}$ | $2.17 \times 10^{-13}$ |
| ZnSO<sub>4</sub> | $7.03 \times 10^{-17}$ | $3.40 \times 10^{-15}$ | $7.24 \times 10^{-15}$ | $1.68 \times 10^{-14}$ | $2.96 \times 10^{-14}$ |
| ZnCO<sub>3</sub> | $5.03 \times 10^{-17}$ | $2.43 \times 10^{-15}$ | $5.17 \times 10^{-15}$ | $1.20 \times 10^{-14}$ | $2.11 \times 10^{-14}$ |
| ZnCl<sup>+</sup> | $2.21 \times 10^{-17}$ | $1.08 \times 10^{-15}$ | $2.31 \times 10^{-15}$ | $5.43 \times 10^{-15}$ | $9.71 \times 10^{-15}$ |
| ZnOH<sup>+</sup> | $2.02 \times 10^{-17}$ | $9.74 \times 10^{-16}$ | $2.07 \times 10^{-15}$ | $4.81 \times 10^{-15}$ | $8.48 \times 10^{-15}$ |
| ZnHCO<sub>3</sub><sup>+</sup> | $1.72 \times 10^{-17}$ | $8.30 \times 10^{-16}$ | $1.77 \times 10^{-15}$ | $4.10 \times 10^{-15}$ | $7.22 \times 10^{-15}$ |
| ZnOHCl | $1.08 \times 10^{-17}$ | $5.25 \times 10^{-16}$ | $1.13 \times 10^{-15}$ | $2.65 \times 10^{-15}$ | $4.73 \times 10^{-15}$ |
| Zn(NTA)<sub>2</sub><sup>4-</sup> | $5.51 \times 10^{-18}$ | $2.50 \times 10^{-16}$ | $5.32 \times 10^{-16}$ | $1.24 \times 10^{-15}$ | $2.18 \times 10^{-15}$ |
| Zn(SO<sub>4</sub>)<sub>2</sub><sup>2-</sup> | $4.36 \times 10^{-18}$ | $2.11 \times 10^{-16}$ | $4.49 \times 10^{-16}$ | $1.04 \times 10^{-15}$ | $1.84 \times 10^{-15}$ |
| ZnCl<sub>2</sub> | $1.14 \times 10^{-18}$ | $5.60 \times 10^{-17}$ | $1.21 \times 10^{-16}$ | $2.88 \times 10^{-16}$ | $5.22 \times 10^{-16}$ |
| Zn(OH)<sub>2</sub> | $7.59 \times 10^{-19}$ | $3.67 \times 10^{-17}$ | $7.82 \times 10^{-17}$ | $1.81 \times 10^{-16}$ | $3.20 \times 10^{-16}$ |
| ZnCl<sub>3</sub><sup>-</sup> | $4.16 \times 10^{-20}$ | $2.06 \times 10^{-18}$ | $4.47 \times 10^{-18}$ | $1.08 \times 10^{-17}$ | $1.98 \times 10^{-17}$ |
| Zn(OH)<sub>3</sub><sup>-</sup> | $2.80 \times 10^{-21}$ | $7.66 \times 10^{-20}$ | $2.88 \times 10^{-19}$ | $6.68 \times 10^{-19}$ | $1.18 \times 10^{-18}$ |
| ZnCl<sub>4</sub><sup>2-</sup> | $1.54 \times 10^{-21}$ | $7.66 \times 10^{-20}$ | $1.68 \times 10^{-19}$ | $4.11 \times 10^{-19}$ | $7.64 \times 10^{-19}$ |
| Zn(OH)<sub>4</sub><sup>2-</sup> | $1.75 \times 10^{-25}$ | $8.44 \times 10^{-24}$ | $1.80 \times 10^{-23}$ | $4.17 \times 10^{-23}$ | $7.35 \times 10^{-23}$ |
| *Total:* | $3.49 \times 10^{-6}$ | $1.57 \times 10^{-4}$ | $3.11 \times 10^{-4}$ | $6.18 \times 10^{-4}$ | $9.26 \times 10^{-4}$ |

**Table S2.** SI values of predicted Zn minerals at different levels of Zn amendment and non-amended controls

| Mineral phase | Chemical formula | Zn amendment level (mg L<sup>-1</sup>) |  |  |  |  |
| --- | --- | --- | --- | --- | --- | --- |
|  |  | 0 | 10 | 20 | 40 | 60 |
| Bianchite | ZnSO<sub>4</sub>.6H<sub>2</sub>O | -16.75 | -15.07 | -14.74 | -14.37 | -14.13 |
| Goslarite | ZnSO<sub>4</sub>.7H<sub>2</sub>O | -16.55 | -14.86 | -14.54 | -14.17 | -13.93 |
| Smithsonite | ZnCO<sub>3</sub> | -11.01 | -9.33 | -9.00 | -8.64 | -8.39 |
| Sphalerite | ZnS | 1.09 | 2.76 | 3.07 | 3.41 | 3.62 |
| Wurtzite | ZnS | -1.38 | 0.29 | 0.60 | 0.93 | 1.15 |
| Zincite | ZnO | -11.40 | -9.72 | -9.39 | -9.02 | -8.78 |
| Zincosite | ZnSO<sub>4</sub> | -22.20 | -20.52 | -20.19 | -19.82 | -19.58 |
| Zn(BO<sub>2</sub>)<sub>2</sub> | Zn(BO<sub>2</sub>)<sub>2</sub> | -22.89 | -21.21 | -20.88 | -20.51 | -20.27 |
| Zn(OH)<sub>2</sub> | Zn(OH)<sub>2</sub> | -12.53 | -10.84 | -10.51 | -10.15 | -9.90 |
| Zn(OH)<sub>2</sub> (am) | Zn(OH)<sub>2</sub> | -12.57 | -10.88 | -10.55 | -10.19 | -9.94 |
| Zn(OH)<sub>2</sub> (beta) | Zn(OH)<sub>2</sub> | -11.84 | -10.16 | -9.83 | -9.46 | -9.22 |
| Zn(OH)<sub>2</sub> (epsilon) | Zn(OH)<sub>2</sub> | -11.62 | -9.94 | -9.61 | -9.25 | -9.00 |
| Zn(OH)<sub>2</sub> (gamma) | Zn(OH)<sub>2</sub> | -12.06 | -10.38 | -10.05 | -9.68 | -9.44 |
| Zn<sub>2</sub>(OH)<sub>2</sub>SO<sub>4</sub> | Zn<sub>2</sub>(OH)<sub>2</sub>SO<sub>4</sub> | -26.34 | -22.97 | -22.31 | -21.58 | -21.09 |
| Zn<sub>2</sub>(OH)<sub>3</sub>Cl | Zn<sub>2</sub>(OH)<sub>3</sub>Cl | -25.00 | -21.63 | -20.97 | -20.24 | -19.74 |
| Zn<sub>3</sub>(PO<sub>4</sub>)<sub>2</sub>.4H<sub>2</sub>O | Zn<sub>3</sub>(PO<sub>4</sub>)<sub>2</sub>.4H<sub>2</sub>O | -30.10 | -25.05 | -24.06 | -22.97 | -22.23 |
| Zn<sub>3</sub>O(SO<sub>4</sub>)<sub>2</sub> | Zn<sub>3</sub>O(SO<sub>4</sub>)<sub>2</sub> | -55.51 | -50.46 | -49.48 | -48.38 | -47.64 |
| Zn<sub>4</sub>(OH)<sub>6</sub>SO<sub>4</sub> | Zn<sub>4</sub>(OH)<sub>6</sub>SO<sub>4</sub> | -47.89 | -41.15 | -39.84 | -38.38 | -37.39 |
| Zn<sub>5</sub>(OH)<sub>8</sub>Cl<sub>2</sub> | Zn<sub>5</sub>(OH)<sub>8</sub>Cl<sub>2</sub> | -58.45 | -50.02 | -48.38 | -46.54 | -45.30 |
| ZnCl<sub>2</sub> | ZnCl<sub>2</sub> | -25.49 | -23.80 | -23.47 | -23.09 | -22.83 |
| ZnCO<sub>3</sub>.H<sub>2</sub>O | ZnCO<sub>3</sub>.H<sub>2</sub>O | -10.80 | -9.12 | -8.79 | -8.42 | -8.18 |
| Zn metal | Zn | -49.07 | -47.38 | -47.06 | -46.69 | -46.44 |
| ZnMoO<sub>4</sub> | ZnMoO<sub>4</sub> | -11.95 | -10.27 | -9.94 | -9.57 | -9.33 |
| ZnO (active) | ZnO | -11.26 | -9.57 | -9.24 | -8.88 | -8.63 |
| ZnS (am) | ZnS | -1.27 | 0.40 | 0.72 | 1.05 | 1.27 |
| ZnSO<sub>4</sub>.H<sub>2</sub>O | ZnSO<sub>4</sub>.H<sub>2</sub>O | -17.75 | -16.06 | -15.73 | -15.37 | -15.12 |

**Table S3.** Distribution (molality) of predicted Cu species at different levels of Cu amendment and non-amended controls

| Valence | Cu species | Cu amendment level (mg L<sup>-1</sup>) |  |  |  |  |
| --- | --- | --- | --- | --- | --- | --- |
|  |  | 0 | 0.5 | 1.5 | 2.5 | 5 |
| Cu(I) | Cu(S4)<sub>2</sub><sup>3-</sup> | $1.16 \times 10^{-12}$ | $1.16 \times 10^{-11}$ | $3.26 \times 10^{-11}$ | $5.38 \times 10^{-11}$ | $1.08 \times 10^{-10}$ |
| | CuS4S5<sup>3-</sup> | $2.05 \times 10^{-13}$ | $2.04 \times 10^{-12}$ | $5.73 \times 10^{-12}$ | $9.46 \times 10^{-12}$ | $1.89 \times 10^{-11}$ |
| | CuCl<sub>2</sub><sup>-</sup> | $8.87 \times 10^{-25}$ | $8.89 \times 10^{-24}$ | $2.53 \times 10^{-23}$ | $4.21 \times 10^{-23}$ | $8.66 \times 10^{-23}$ |
| | CuCl | $9.56 \times 10^{-26}$ | $9.57 \times 10^{-25}$ | $2.72 \times 10^{-24}$ | $4.53 \times 10^{-24}$ | $9.30 \times 10^{-24}$ |
| | CuCl<sub>3</sub><sup>2-</sup> | $1.38 \times 10^{-26}$ | $1.38 \times 10^{-25}$ | $3.92 \times 10^{-25}$ | $6.55 \times 10^{-25}$ | $1.35 \times 10^{-24}$ |
| | Cu<sup>+</sup> | $3.22 \times 10^{-27}$ | $3.22 \times 10^{-26}$ | $9.14 \times 10^{-26}$ | $1.52 \times 10^{-25}$ | $3.12 \times 10^{-25}$ |
| Cu(II) | Cu(HS)<sub>3</sub><sup>-</sup> | $8.84 \times 10^{-7}$ | $8.79 \times 10^{-6}$ | $2.46 \times 10^{-5}$ | $4.04 \times 10^{-5}$ | $8.00 \times 10^{-5}$ |
| | Cu(NTA)<sup>-</sup> | $4.71 \times 10^{-21}$ | $4.72 \times 10^{-20}$ | $1.34 \times 10^{-19}$ | $2.23 \times 10^{-19}$ | $4.57 \times 10^{-19}$ |
| | CuOH(NTA)<sup>2-</sup> | $2.30 \times 10^{-22}$ | $2.30 \times 10^{-21}$ | $6.54 \times 10^{-21}$ | $1.09 \times 10^{-20}$ | $2.23 \times 10^{-20}$ |
| | Cu(NTA)<sub>2</sub><sup>4-</sup> | $1.79 \times 10^{-24}$ | $1.79 \times 10^{-23}$ | $5.09 \times 10^{-23}$ | $8.48 \times 10^{-23}$ | $1.74 \times 10^{-22}$ |
| | CuCO<sub>3</sub> | $1.17 \times 10^{-24}$ | $1.17 \times 10^{-23}$ | $3.32 \times 10^{-23}$ | $5.53 \times 10^{-23}$ | $1.13 \times 10^{-22}$ |
| | CuNH<sub>3</sub><sup>2+</sup> | $2.59 \times 10^{-25}$ | $2.60 \times 10^{-24}$ | $7.36 \times 10^{-24}$ | $1.23 \times 10^{-23}$ | $2.51 \times 10^{-23}$ |
| | Cu<sup>+2</sup> | $1.17 \times 10^{-25}$ | $1.17 \times 10^{-24}$ | $3.33 \times 10^{-24}$ | $5.54 \times 10^{-24}$ | $1.14 \times 10^{-23}$ |
| | CuOH<sup>+</sup> | $1.13 \times 10^{-25}$ | $1.13 \times 10^{-24}$ | $3.22 \times 10^{-24}$ | $5.36 \times 10^{-24}$ | $1.10 \times 10^{-23}$ |
| | Cu(CO<sub>3</sub>)<sub>2</sub><sup>2-</sup> | $6.61 \times 10^{-26}$ | $6.61 \times 10^{-25}$ | $1.88 \times 10^{-24}$ | $3.13 \times 10^{-24}$ | $6.40 \times 10^{-24}$ |
| | CuSO<sub>4</sub> | $1.70 \times 10^{-26}$ | $1.70 \times 10^{-25}$ | $4.83 \times 10^{-25}$ | $8.05 \times 10^{-25}$ | $1.65 \times 10^{-24}$ |
| | CuHCO<sub>3</sub><sup>+</sup> | $7.78 \times 10^{-27}$ | $7.79 \times 10^{-26}$ | $2.21 \times 10^{-25}$ | $3.68 \times 10^{-25}$ | $7.55 \times 10^{-25}$ |
| | Cu(OH)<sub>2</sub> | $6.87 \times 10^{-27}$ | $6.88 \times 10^{-26}$ | $1.95 \times 10^{-25}$ | $3.25 \times 10^{-25}$ | $6.66 \times 10^{-25}$ |
| | CuH(NTA) | $4.28 \times 10^{-27}$ | $4.29 \times 10^{-26}$ | $1.22 \times 10^{-25}$ | $2.03 \times 10^{-25}$ | $4.15 \times 10^{-25}$ |
| | CuCl<sup>+</sup> | $3.24 \times 10^{-27}$ | $3.24 \times 10^{-26}$ | $9.21 \times 10^{-26}$ | $1.53 \times 10^{-25}$ | $3.15 \times 10^{-25}$ |
| | CuCl<sub>2</sub> | $3.76 \times 10^{-29}$ | $3.77 \times 10^{-28}$ | $1.07 \times 10^{-27}$ | $1.79 \times 10^{-27}$ | $3.67 \times 10^{-27}$ |
| | Cu(OH)<sub>3</sub><sup>-</sup> | $1.04 \times 10^{-29}$ | $1.04 \times 10^{-28}$ | $2.94 \times 10^{-28}$ | $4.90 \times 10^{-28}$ | $1.00 \times 10^{-27}$ |
| | CuCl<sub>3</sub><sup>-</sup> | $1.72 \times 10^{-32}$ | $1.73 \times 10^{-31}$ | $4.91 \times 10^{-31}$ | $8.19 \times 10^{-31}$ | $1.69 \times 10^{-30}$ |
| | Cu(OH)<sub>4</sub><sup>2-</sup> | $1.28 \times 10^{-34}$ | $1.28 \times 10^{-33}$ | $3.63 \times 10^{-33}$ | $6.05 \times 10^{-33}$ | $1.24 \times 10^{-32}$ |
| | CuCl<sub>4</sub><sup>2-</sup> | $5.21 \times 10^{-36}$ | $5.22 \times 10^{-35}$ | $1.49 \times 10^{-34}$ | $2.48 \times 10^{-34}$ | $5.12 \times 10^{-34}$ |
| | *Total:* | $8.84 \times 10^{-7}$ | $8.79 \times 10^{-6}$ | $2.46 \times 10^{-5}$ | $4.04 \times 10^{-5}$ | $8.00 \times 10^{-5}$ |

**Table S4.**
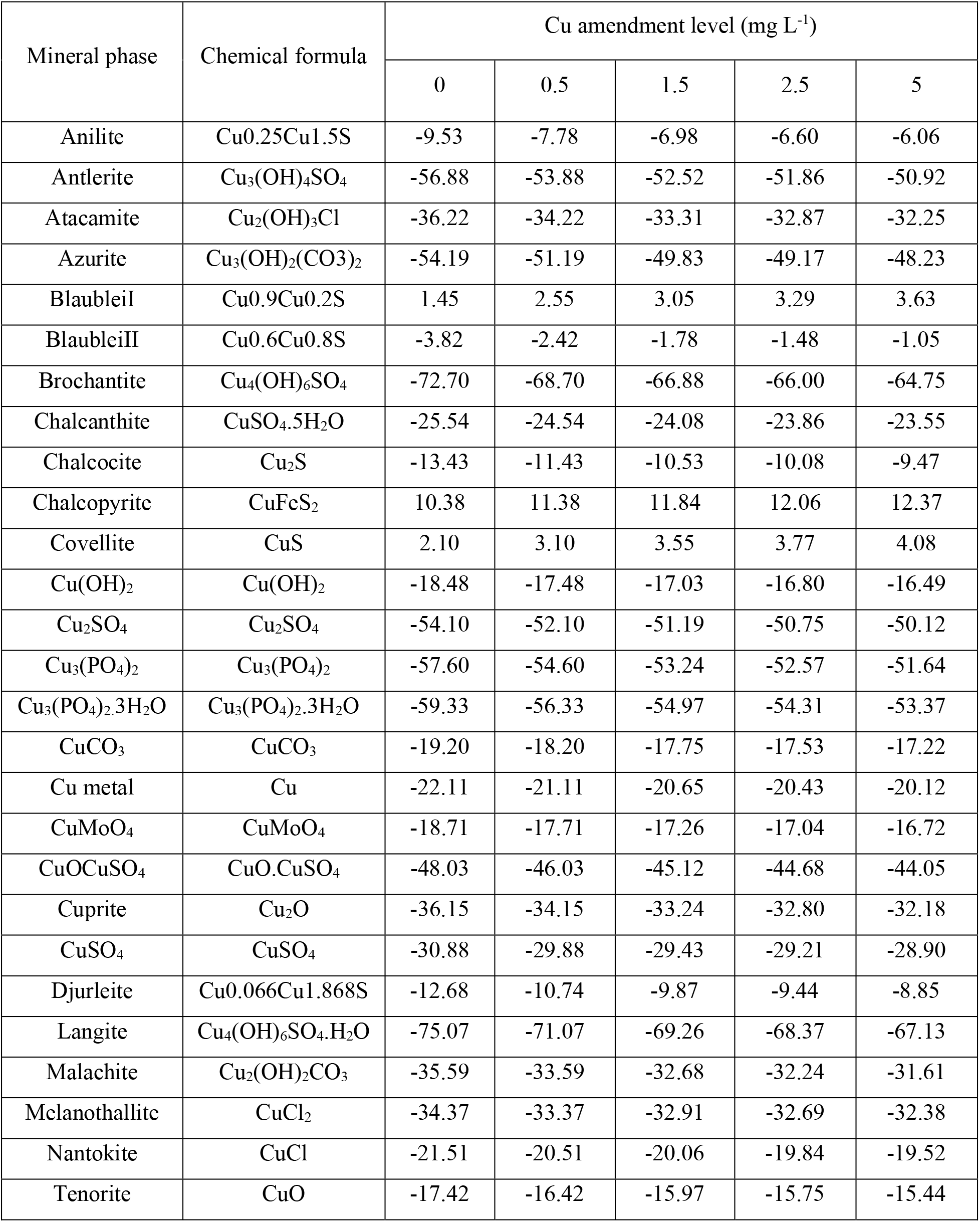
SI values of predicted Cu minerals at different levels of Cu amendment and non-amended controls

**Table S5.**
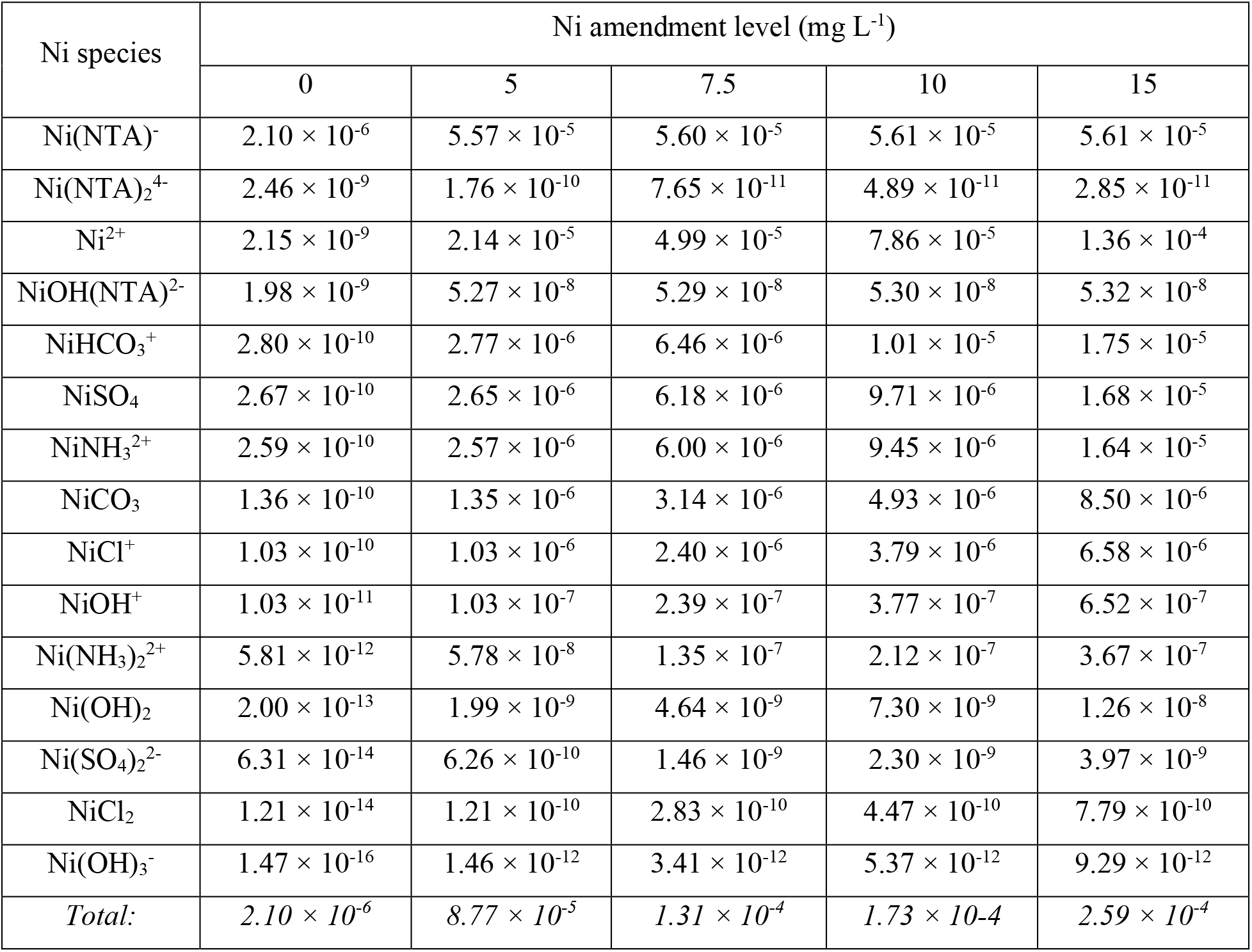
Distribution (molality) of predicted Ni species at different levels of Ni amendment and non-amended controls

**Table S6.** SI values of predicted Ni minerals at different levels of Ni amendment and non-amended controls

| Mineral phase | Chemical formula | Ni amendment level (mg L<sup>-1</sup>) |  |  |  |  |
| --- | --- | --- | --- | --- | --- | --- |
|  |  | 0 | 5 | 7.5 | 10 | 15 |
| Bunsenite | NiO | -5.86 | -1.86 | -1.49 | -1.30 | -1.06 |
| Morenosite | NiSO<sub>4</sub>·7H<sub>2</sub>O | -9.79 | -5.79 | -5.42 | -5.23 | -4.99 |
| Ni(OH)<sub>2</sub> | Ni(OH)<sub>2</sub> | -6.22 | -2.22 | -1.86 | -1.66 | -1.42 |
| Ni<sub>3</sub>(PO<sub>4</sub>)<sub>2</sub> | Ni<sub>3</sub>(PO<sub>4</sub>)<sub>2</sub> | -14.35 | -2.36 | -1.26 | -0.67 | 0.04 |
| Ni<sub>4</sub>(OH)<sub>6</sub>SO<sub>4</sub> | Ni<sub>4</sub>(OH)<sub>6</sub>SO<sub>4</sub> | -25.00 | -9.02 | -7.54 | -6.76 | -5.81 |
| NiCO<sub>3</sub> | NiCO<sub>3</sub> | -7.45 | -3.45 | -3.08 | -2.89 | -2.65 |
| NiMoO<sub>4</sub> | NiMoO<sub>4</sub> | -4.35 | -0.35 | 0.01 | 0.21 | 0.45 |
| NiS (alpha) | NiS | 1.95 | 5.94 | 6.31 | 6.51 | 6.75 |
| NiS (beta) | NiS | 7.45 | 11.44 | 11.81 | 12.01 | 12.25 |
| NiS (gamma) | NiS | 9.15 | 13.14 | 13.51 | 13.71 | 13.95 |
| Retgersite | NiSO<sub>4</sub>·6H<sub>2</sub>O | -9.87 | -5.87 | -5.51 | -5.31 | -5.07 |

**Table S7.**
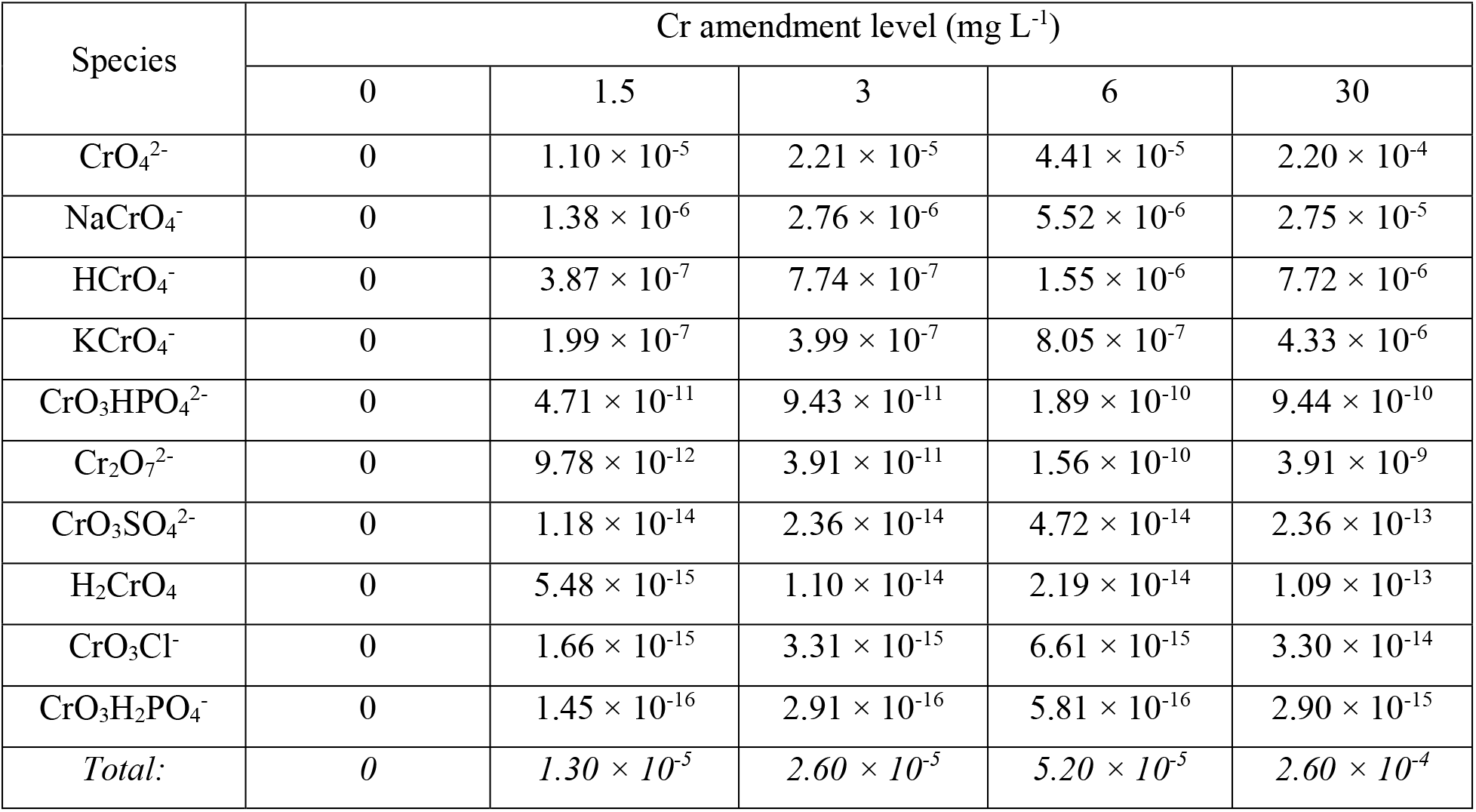
Distribution (molality) of predicted Cr species at different levels of Cr amendment and non-amended controls

**Table S8.** SI values of predicted Cr minerals at different levels of Cr amendment and non-amended controls

| Mineral phase | Chemical formula | Cr amendment level (mg L<sup>-1</sup>) |  |  |  |  |
| --- | --- | --- | --- | --- | --- | --- |
|  |  | 0* | 1.5 | 3 | 6 | 30 |
| (NH<sub>4</sub>)<sub>2</sub>CrO<sub>4</sub> | (NH<sub>4</sub>)<sub>2</sub>CrO<sub>4</sub> | - | -10.09 | -9.79 | -9.49 | -8.80 |
| CaCrO<sub>4</sub> | CaCrO<sub>4</sub> | - | -5.62 | -5.32 | -5.02 | -4.33 |
| CrO<sub>3</sub> | CrO<sub>3</sub> | - | -17.57 | -17.27 | -16.96 | -16.27 |
| CuCrO<sub>4</sub> | CuCrO<sub>4</sub> | - | -25.32 | -25.02 | -24.72 | -24.02 |
| K<sub>2</sub>Cr<sub>2</sub>O<sub>7</sub> | K<sub>2</sub>Cr<sub>2</sub>O<sub>7</sub> | - | -13.26 | -12.66 | -12.05 | -10.59 |
| K<sub>2</sub>CrO<sub>4</sub> | K<sub>2</sub>CrO<sub>4</sub> | - | -9.02 | -8.72 | -8.41 | -7.64 |
| MgCrO<sub>4</sub> | MgCrO<sub>4</sub> | - | -13.38 | -13.07 | -12.77 | -12.08 |
| Na<sub>2</sub>Cr<sub>2</sub>O<sub>7</sub> | Na<sub>2</sub>Cr<sub>2</sub>O<sub>7</sub> | - | -19.01 | -18.41 | -17.81 | -16.41 |
| Na<sub>2</sub>CrO<sub>4</sub> | Na<sub>2</sub>CrO<sub>4</sub> | - | -10.92 | -10.62 | -10.32 | -9.62 |

\* None of Cr minerals were predicted in non-amended trials of Cr tolerance experiment

**Table S9.** SI values for predicted As minerals at different levels of As amendment and non-amended controls

| Species | As amendment level (mg L<sup>-1</sup>) |  |  |  |  |
| --- | --- | --- | --- | --- | --- |
|  | 0 | 10 | 30 | 300 | 500 |
| HAsO<sub>4</sub><sup>2-</sup> | 0 | $6.84 \times 10^{-5}$ | $2.05 \times 10^{-4}$ | $2.06 \times 10^{-3}$ | $3.43 \times 10^{-3}$ |
| H<sub>2</sub>AsO<sub>4</sub><sup>-</sup> | 0 | $3.91 \times 10^{-6}$ | $1.17 \times 10^{-5}$ | $1.13 \times 10^{-4}$ | $1.82 \times 10^{-4}$ |
| AsO<sub>4</sub><sup>3-</sup> | 0 | $8.08 \times 10^{-8}$ | $2.44 \times 10^{-7}$ | $2.62 \times 10^{-6}$ | $4.59 \times 10^{-6}$ |
| H<sub>3</sub>AsO<sub>4</sub> | 0 | $9.50 \times 10^{-12}$ | $2.84 \times 10^{-11}$ | $2.69 \times 10^{-10}$ | $4.31 \times 10^{-10}$ |
| *Total:* | 0 | $7.24 \times 10^{-5}$ | $2.17 \times 10^{-4}$ | $2.17 \times 10^{-3}$ | $3.62 \times 10^{-3}$ |

**Table S10.** SI values of predicted As minerals at different levels of As amendment and non-amended controls

| Phase | Chemical formula | As amendment level (mg L<sup>-1</sup>) |  |  |  |  |
| --- | --- | --- | --- | --- | --- | --- |
|  |  | 0 | 10 | 30 | 300 | 500 |
| AlAsO<sub>4</sub>·2H<sub>2</sub>O | AlAsO<sub>4</sub>·2H<sub>2</sub>O | - | -8.71 | -8.24 | -7.26 | -7.06 |
| As<sub>2</sub>O<sub>5</sub> | As<sub>2</sub>O<sub>5</sub> | - | -28.66 | -27.71 | -25.76 | -25.35 |
| Ca<sub>3</sub>(AsO<sub>4</sub>)<sub>2</sub>·4H<sub>2</sub>O | Ca<sub>3</sub>(AsO<sub>4</sub>)<sub>2</sub>·4H<sub>2</sub>O | - | -5.85 | -4.90 | -2.97 | -2.57 |
| Co<sub>3</sub>(AsO<sub>4</sub>)<sub>2</sub> | Co<sub>3</sub>(AsO<sub>4</sub>)<sub>2</sub> | - | -12.21 | -11.26 | -9.33 | -8.93 |
| Cu<sub>3</sub>(AsO<sub>4</sub>)<sub>2</sub>·2H<sub>2</sub>O | Cu<sub>3</sub>(AsO<sub>4</sub>)<sub>2</sub>·2H<sub>2</sub>O | - | -58.02 | -57.07 | -55.07 | -54.62 |
| Mn<sub>3</sub>(AsO<sub>4</sub>)<sub>2</sub>·8H<sub>2</sub>O | Mn<sub>3</sub>(AsO<sub>4</sub>)<sub>2</sub>·8H<sub>2</sub>O | - | -7.25 | -6.31 | -4.42 | -4.05 |
| Ni<sub>3</sub>(AsO<sub>4</sub>)<sub>2</sub>·8H<sub>2</sub>O | Ni<sub>3</sub>(AsO<sub>4</sub>)<sub>2</sub>·8H<sub>2</sub>O | - | -18.84 | -17.89 | -15.96 | -15.56 |
| Zn<sub>3</sub>(AsO<sub>4</sub>)<sub>2</sub>·2.5H<sub>2</sub>O | Zn<sub>3</sub>(AsO<sub>4</sub>)<sub>2</sub>·2.5H<sub>2</sub>O | - | -36.65 | -35.69 | -33.71 | -33.28 |

\* None of As minerals were predicted in non-amended trials of As tolerance experiment

